# RECON infers regions of interest from H&E images and reconstructs whole-slide molecular profiles at single-cell resolution

**DOI:** 10.64898/2026.08.25.747122

**Authors:** Xinxing Yang, Ninghui Hao, Running Zhao, Sofia Angel, Yusheng Tan, Christine G. Lian, Li Zhou, Daniel Olson, Kun-Hsing Yu, Arlene Ruiz de Luzuriaga, Guihong Wan

**Affiliations:** Institute for Population and Precision Health, Department of Family Medicine, The University of Chicago, Chicago, 60637, Illinois, USA; Section of Dermatology, The University of Chicago, Chicago, 60637, Illinois, USA; Department of Pathology, Brigham and Women’s Hospital, Harvard Medical School, Boston, 02115, Massachusetts, USA; Department of Medicine, Brigham and Women’s Hospital, Harvard Medical School, Boston, 02115, Massachusetts, USA; Department of Medicine, The University of Chicago, Chicago, 60637, Illinois, USA; Department of Biomedical Informatics, Harvard Medical School, Boston, 02115, Massachusetts, USA; Harvard Data Science Initiative, Harvard University, Cambridge, 02134, Massachusetts, USA; Kempner Institute for the Study of Natural and Artificial Intelligence, Harvard University, Cambridge, 02134, Massachusetts, USA

**Keywords:** Spatial omics, H&E images, Regions of interest selection, Gene expression prediction, Whole-slide molecular reconstruction

## Abstract

Spatial omics technologies resolve molecular expression and spatial architecture at single-cell resolution, but profiling whole slides remains costly. In practice, only a few regions of interest (ROIs) are profiled, leaving the rest of the tissue unmeasured. S2-omics was the first framework to unify ROI selection with out-of-ROI prediction, but it operates on superpixels rather than individual cells and predicts discrete cell types rather than continuous molecular profiles. Superpixel-based representations do not explicitly preserve cell boundaries, while categorical cell-type labels cannot quantify molecular expression within cells. Here we present RECON, a two-stage framework that performs ROI inference and whole-slide molecular reconstruction at single-cell resolution, predicting both continuous molecular profiles and discrete cell-type labels. In the first stage, RECON extracts morphological and microenvironmental features from individual cells to identify a representative ROI for spatially resolved single-cell molecular profiling. In the second stage, RECON trains deep learning models on molecular measurements acquired within the selected ROI and reconstructs transcriptomic or proteomic profiles for all remaining cells on the slide. Benchmarked against pathologist annotations, RECON’s ROI selection outperforms the superpixel-based S2-omics approaches (IoU: 0.75 versus 0.64). For transcriptomics, refining the modeling unit from superpixels to single cells improves per-gene Pearson correlation by 22%. For proteomics, RECON surpasses the current state-of-the-art method, ROSIE, across all 16 markers, with a median per-cell Pearson correlation of 0.91 versus 0.84. Moreover, RECON delineates tumour boundaries and regions with distinct immune-cell densities, and highlights candidate tertiary lymphoid structures. Together, these results demonstrate that RECON enables informative ROI selection and whole-slide molecular reconstruction at single-cell resolution for both spatial transcriptomics and spatial proteomics.

## Introduction

Spatial omics technologies such as Xenium [1], PhenoCycler [2], and Orion [3] resolve both molecular expression and tissue spatial architecture at single-cell resolution and have become central tools for dissecting the tumor microenvironment [4, 5]. However, profiling an entire tissue section remains costly. For example, reagents and service fees for a Xenium run can amount to several thousand US dollars [6]. In practice, only a few regions of interest (ROIs) are typically measured, leaving the molecular profiles of the remaining tissue unmeasured [7]. At present, ROIs are typically outlined by pathologists on H&E sections. However, manual ROI selection is subjective and may not be reliably reproducible. It also relies on visual morphology alone, yet regions with similar histological appearance can differ markedly in molecular composition [8, 9]. These limitations motivate an automated alternative: a computational approach that selects an informative ROI from the H&E image for spatial molecular profiling and reconstructs molecular profiles across the remainder of the section [10, 11].

S2-omics [7] unifies automated ROI selection and out-of-ROI cell-type prediction. It clusters and scores superpixels on the H&E image, selects the most representative ROI, and then predicts the cell types of the remaining cells outside that ROI. However, S2-omics has two fundamental limitations. First, S2-omics operates on superpixels, which are geometric tiles laid out on a regular grid across the slide. Although their side length can be reduced to 8 *µm*, approaching the scale of a single cell, the tiling is defined without reference to morphology or cell boundaries [12, 13]. Grid lines may split individual cells across several adjacent tiles, and no single tile corresponds to exactly one cell. Therefore, unlike segmented cells [14–17], geometric tiles do not faithfully delineate individual cells and are not biologically meaningful units of analysis. Second, S2-omics predicts discrete cell types rather than continuous molecular expression. Cell-type labels cannot answer quantitative questions, such as how much of a given gene or protein a cell expresses [18–20]. In short, S2-omics serves as the first framework to unify ROI selection and out-of-ROI prediction, but it neither operates on segmented cells nor produces continuous molecular profiles at the cellular level.

Beyond S2-omics, related methods [21–29] generally address only one of the two stages, each with key limitations. For ROI selection, automated approaches remain scarce. ROIs are typically delineated manually by pathologists, with S2-omics being, to date, the only framework that attempts to automate this step. In contrast, predicting molecular information from H&E images has attracted considerable attention, but these methods focus on molecular prediction alone and do not address ROI selection. In spatial transcriptomics, iSCALE [26] extrapolates gene expression at super-resolution by integrating multiple sections, and GHIST [27] was the first to predict gene expression using segmented cells as the modeling unit. In spatial proteomics, ROSIE [28] predicts multiplexed protein expression at the pixel level, and GigaTIME [29] translates H&E images into pixel-level virtual multiplexed immunofluorescence (mIF) images. More fundamentally, these methods are trained under whole-slide supervision, assuming molecular ground truth is available across the whole slide. This assumption fails in ROI-driven settings, where typically only a single ROI is profiled per slide. Taken together, no existing method addresses both ROI selection and molecular prediction at single-cell resolution, supports extrapolation from in-ROI measurements, and spans transcriptomics and proteomics.

To address these gaps, we present RECON, a two-stage framework that infers an ROI from the H&E image and reconstructs whole-slide molecular profiles at single-cell resolution. RECON operates on segmented cells as the basic unit of analysis at both stages, rather than superpixels. It first extracts morphological and microenvironmental features for each cell and groups cells into functional clusters, from which a scoring function selects the most representative ROI among candidate regions. Then, the spatially resolved molecular measurements acquired within the selected ROI serve as training data to reconstruct molecular profiles for all cells in the remaining tissue. Both stages share a single UNI backbone [30] and a common per-cell feature-extraction pipeline, allowing the same architecture to be applied to both spatial transcriptomics and spatial proteomics. Thus, RECON simultaneously satisfies three requirements not previously met in one framework: single-cell resolution, molecular extrapolation from a profiled ROI, and cross-modality applicability.

We validated RECON across colorectal, breast, and gastric cancers, spanning both transcriptomic and proteomic modalities. In colorectal cancer (CRC) sections, the ROIs inferred by RECON agreed more closely with expert annotations than those inferred by superpixel-level S2-omics (IoU: 0.75 versus 0.64). By representing tissue at single-cell resolution, RECON preserved tissue-compartment boundaries that were blurred by superpixel averaging. In breast cancer Xenium sections, RECON reconstructed transcriptomic expression at single-cell resolution and outperformed the cell-level GHIST [27]. The benefits of single-cell resolution extended beyond quantitative accuracy. In a gastric cancer Xenium section, RECON recovered tertiary lymphoid structures, tumor boundaries smaller than a single superpixel, and regions with distinct immune densities. On a separate CRC Orion dataset, the same architecture used protein expression within the ROI as training data to reconstruct protein expression for all cells outside the ROI, outperforming the pixel-level ROSIE [28] across all 16 protein markers. Ablation experiments further showed that refining the modeling unit from superpixels to single cells improved per-gene Pearson correlation by 22%.

## Results

### Overview of the RECON framework

Fig. 1 presents an overview of the RECON framework. RECON uses two adjacent sections from the same FFPE block to perform automated ROI selection and whole-slide molecular reconstruction at single-cell resolution (Fig. 1a). The first section is stained with H&E to infer an informative ROI. Because adjacent sections from the same block are histologically concordant, the selected ROI is registered to the second section for within-ROI spatial omics profiling. These molecular measurements then serve as training data to reconstruct molecular profiles for all cells outside the ROI.

**Fig. 1.**
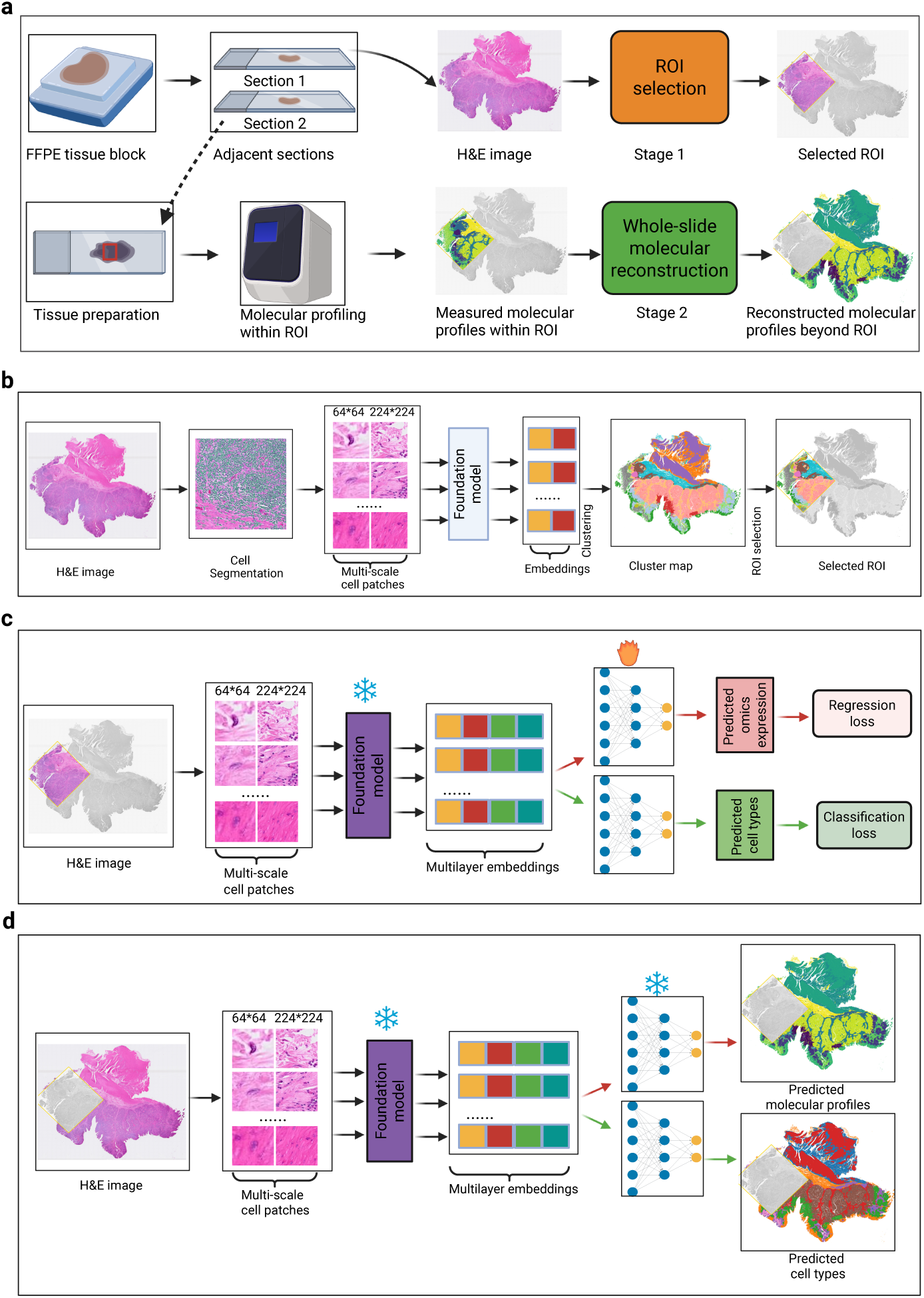
Overview of the RECON framework. **a**, RECON operates on two adjacent sections from the same FFPE block, one imaged by H&E and the other profiled by spatial omics. Stage 1 infers an ROI from the H&E image; Stage 2 reconstructs molecular profiles for all cells outside the ROI. **b**, Stage 1. Cells are segmented from the H&E image and encoded by a frozen pathology foundation model at two scales. Cell embeddings are clustered into a histological map, from which candidate windows are scored for representativeness and the highest-scoring window is selected as the ROI. **c**, Stage 2. Molecular measurements within the ROI supervise a lightweight prediction head with two independent branches: one regressing continuous molecular expression and the other classifying cell types. Both branches operate on the same per-cell morphological features. **d**, Output. Each cell outside the ROI receives a continuous molecular profile and a cell-type assignment, yielding a whole-slide single-cell map from a routine H&E section.

In Stage 1 (Fig. 1b), RECON segments cells on the H&E section and crops multi-scale patches centered on each segmented cell (64 × 64 and 224 × 224 pixels). A frozen pathology foundation model (e.g., UNI [30]) encodes these patches into multi-scale cell embeddings, which are clustered to yield a histological map of the slide. RECON then scores candidate windows according to how closely their cluster composition matches that of the whole slide and selects the highest-scoring window as the ROI. In Stage 2 (Fig. 1c), RECON reconstructs molecular profiles of cells outside the ROI, training solely on the measurements acquired within the ROI. It reuses the same cell masks, patches, and frozen encoder, and trains only a lightweight prediction head on top. The prediction head has two branches: a regression branch that reconstructs continuous molecular expression by regression loss, and a classification branch that assigns cell types by classification loss. Once trained, the prediction head is applied to all remaining cells, yielding the continuous molecular expression and cell-type assignments to individual cells on the slide (Fig. 1d). Both stages of RECON run without modification on spatial transcriptomics and spatial proteomics.

### Single-cell-level ROI selection outperforms superpixel-based approaches

We first asked whether RECON’s ROI selection agrees with expert annotation. To answer this question, we evaluated different ROI sizes (6.5, 3, and 2 mm per side) using two neighboring sections from a human CRC FFPE block [31], one profiled by Visium HD and the other by Xenium. Each section had a corresponding expert ROI annotation (Fig. 2a). The two corresponding H&E images were aligned using global affine registration based on SIFT feature matching [32] (Fig. 2a, bottom; green lines denote matched keypoint pairs). ROI agreement was quantified by Intersection over Union (IoU) between the selected ROI and the expert annotation.

**Fig. 2.**
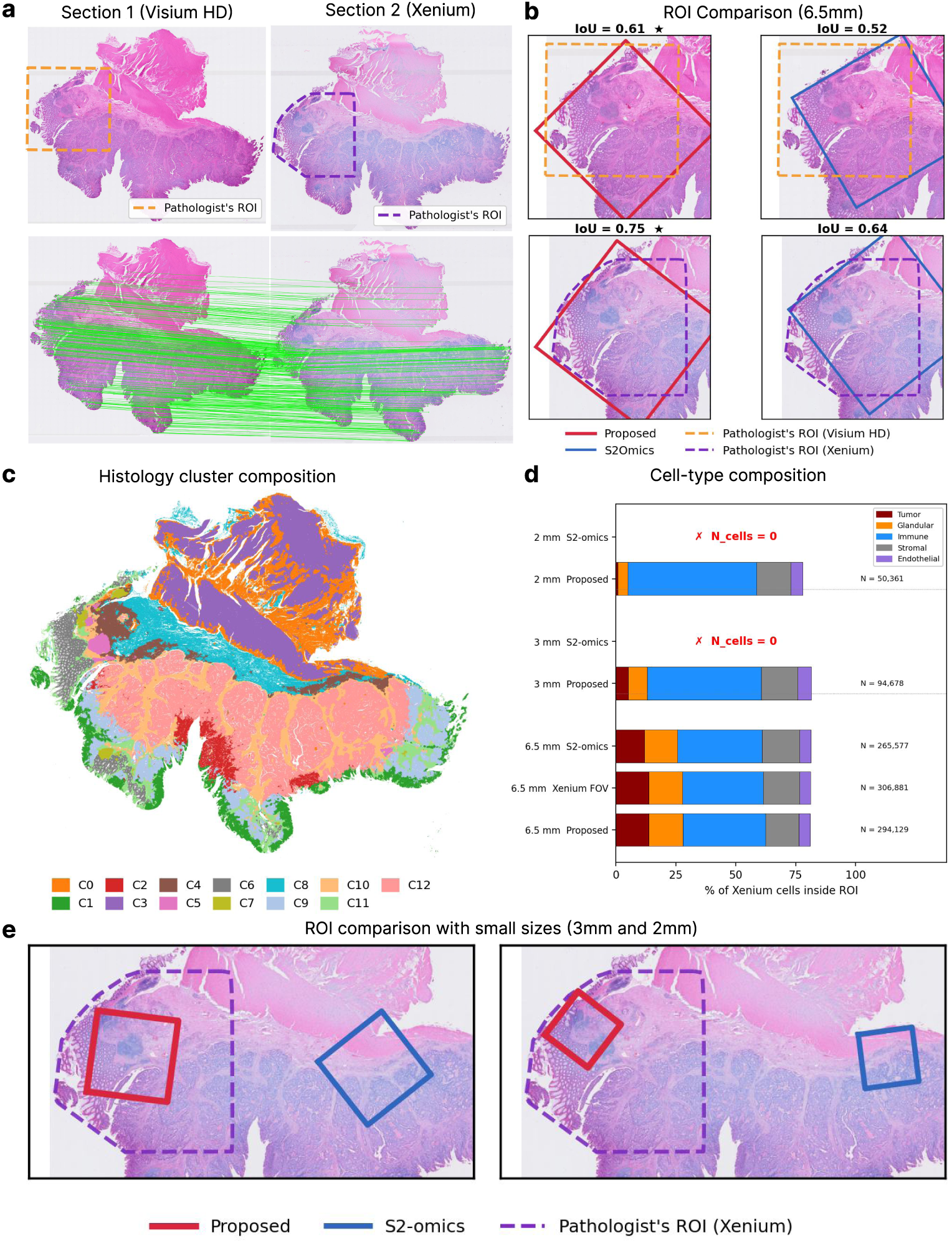
RECON infers ROIs that agree with expert annotations. **a**, Adjacent sections profiled by Visium HD and Xenium, each with a pathologist annotation; and the keypoints used for registration. **b**, ROIs selected by RECON and S2-omics at 6.5 mm against pathologist annotations on the Visium HD (top) and Xenium (bottom) sections, with Intersection over Union (IoU) above each panel. **c**, Histological cluster map used for ROI scoring. **d**, Cell-type composition and number of Xenium cells captured within each selected ROI at 2, 3 and 6.5 mm. **e**, At smaller sizes, ROIs selected by S2-omics drifted beyond the Xenium field of view (FOV), whereas ROIs inferred by RECON remained within the annotated region.

RECON’s ROIs (6.5 mm) agreed more closely with expert annotations than those of superpixel-based S2-omics for both sections: IoU 0.61 versus 0.52 on Visium HD and 0.75 versus 0.64 on Xenium (Fig. 2b). This advantage stems from the choice of computational unit. Before ROI selection, RECON clusters segmented cells and organizes the whole section into 13 histological clusters (Fig. 2c), preserving sharp boundaries between compartments such as tumor, glands, and stroma. The scoring function then selects the most representative region on this basis. In contrast, superpixel-based methods average within regular square tiles, which can blur boundaries between adjacent compartments and shift the selected ROI away from the expert annotation.

The difference was amplified at smaller ROI sizes (Fig. 2d, e), where reducing the profiled area offers greater cost savings and accurate ROI selection is therefore particularly important. N cells refer to the number of cells within the selected ROI actually profiled by Xenium, which determines whether the ROI can provide enough training data for downstream reconstruction. At ROI sizes of 2 and 3 mm, ROIs selected by S2-omics departed from the expert annotations and even drifted outside the Xenium field of view (FOV) (Fig. 2e), containing no profiled cells and therefore providing no molecular measurements for training (N cells = 0; Fig. 2d). By comparison, RECON’s ROIs at these sizes fell entirely within the expert annotation, containing 50,361 and 94,678 cells at 2 and 3 mm, respectively. The advantage persisted at 6.5 mm: RECON’s ROI contained 294,129 cells versus 265,577 for S2-omics, closer to the 306,881 cells of the Xenium FOV, and its cell-type composition matched more closely than that of the Xenium FOV (Fig. 2d). Together, these results indicate that RECON-selected ROIs recapitulate pathologist annotations and capture cellular composition representative of the whole section.

### RECON infers ROIs and reconstructs spatial transcriptomes at single-cell resolution

Because whole-slide single-cell transcriptomic profiling remains expensive, we next asked whether molecular profiles outside the selected ROI could be reconstructed from measurements within the ROI. We evaluated the full two-stage RECON framework on two human breast cancer Xenium sections: one profiled by Xenium V1 with a 313-gene panel [33], and the other by Xenium Prime with a 5,000-gene panel, from which the 300 most variable genes served as reconstruction targets. For each section, RECON selected an ROI from the H&E image and used only the molecular measurements within the selected ROI for training and reconstructed single-cell transcriptomes for all cells outside the ROI, with their measured profiles held out for evaluation.

RECON selected ROIs using clusters of segmented cells derived from H&E features (H&E ROIs). We compared these H&E ROIs with ROIs obtained by clustering cells based on their measured gene expression (Gene ROIs). Reconstruction accuracy was assessed using Pearson correlation for models trained on either H&E ROI data or Gene ROI data, and was also compared with the single-cell GHIST transcriptome prediction method [27]. To further assess the generalization of our framework, we compared reconstruction accuracy as a function of distance from a selected ROI.

The H&E and Gene ROIs overlapped for both sections (IoU: 0.631 and 0.404; Fig. 3a, b). Notably, using the respective measurements from the H&E and Gene ROIs yielded comparable molecular reconstruction. When evaluated on cells outside both ROIs, per-cell Pearson correlations were nearly identical (0.632 versus 0.632 for the V1 section; 0.222 versus 0.227 for the Prime section; Supplementary Figs. 1, 2). The H&E ROIs were also enriched for tumor-associated clusters (Fig. 3a,b, inner versus outer rings). For the first section (Fig. 3a), the dominant tumor cluster C5 accounted for 34.2% of cells in the whole section versus 44.8% of cells within the ROI.

**Fig. 3.**
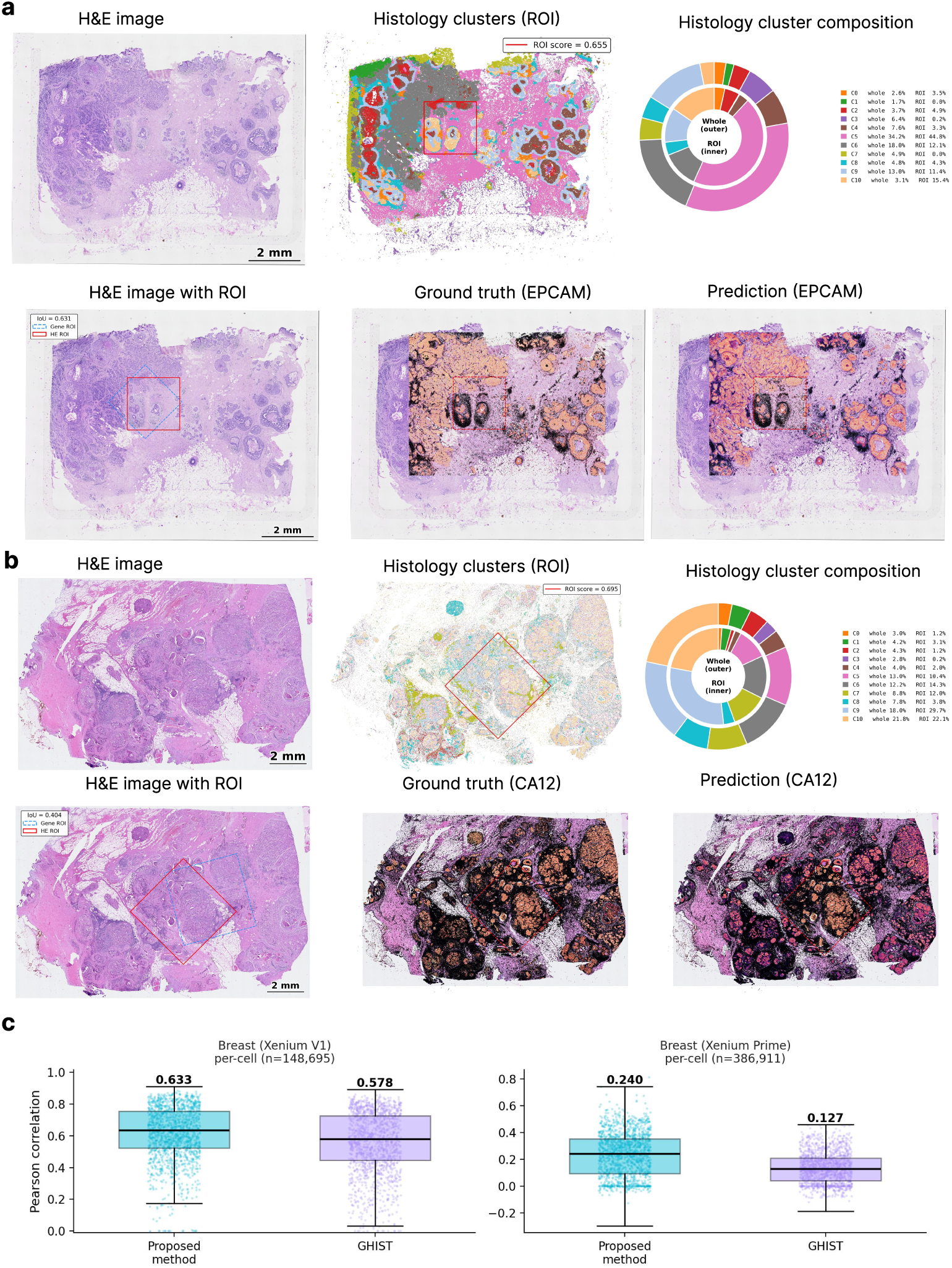
RECON infers ROIs and reconstructs spatial transcriptomes at single-cell resolution. **a**, Xenium V1 breast cancer section. Top: H&E image, histological cluster map with the selected ROI, and cluster composition within the ROI (inner ring) compared with the whole section (outer ring). Bottom: the H&E-inferred ROI (HE ROI) against the ROI obtained from measured gene expression (Gene ROI), and measured versus reconstructed expression of EPCAM. **b**, As in **a**, for the Xenium Prime breast cancer section, with CA12 shown as the representative gene. **c**, Per-cell Pearson correlation between measured and reconstructed expression for RECON and GHIST, computed on cells outside the ROI. Boxes show the median and interquartile range; whiskers extend to 1.5 times the interquartile range.

The spatial distributions of predicted marker genes showed close agreement with the ground truth. The reconstructed EPCAM pattern reproduced the glandular and epithelial architecture in the V1 section (Fig. 3a), and the reconstructed CA12 delineated tumor regions consistent with the measured data in the Prime section (Fig. 3b). Other epithelial and tumor markers showed similar spatial concordance for both Xenium platforms. Per-gene Pearson correlations exceeded 0.72 for KRT7, KRT8, and FASN for the V1 data (Supplementary Figs. 1c, 3a) and ranged from 0.58 to 0.70 for CA12, XBP1, and RAB11FIP1 for the Prime data (Supplementary Figs. 2c, 4a).

**Fig. 4.**
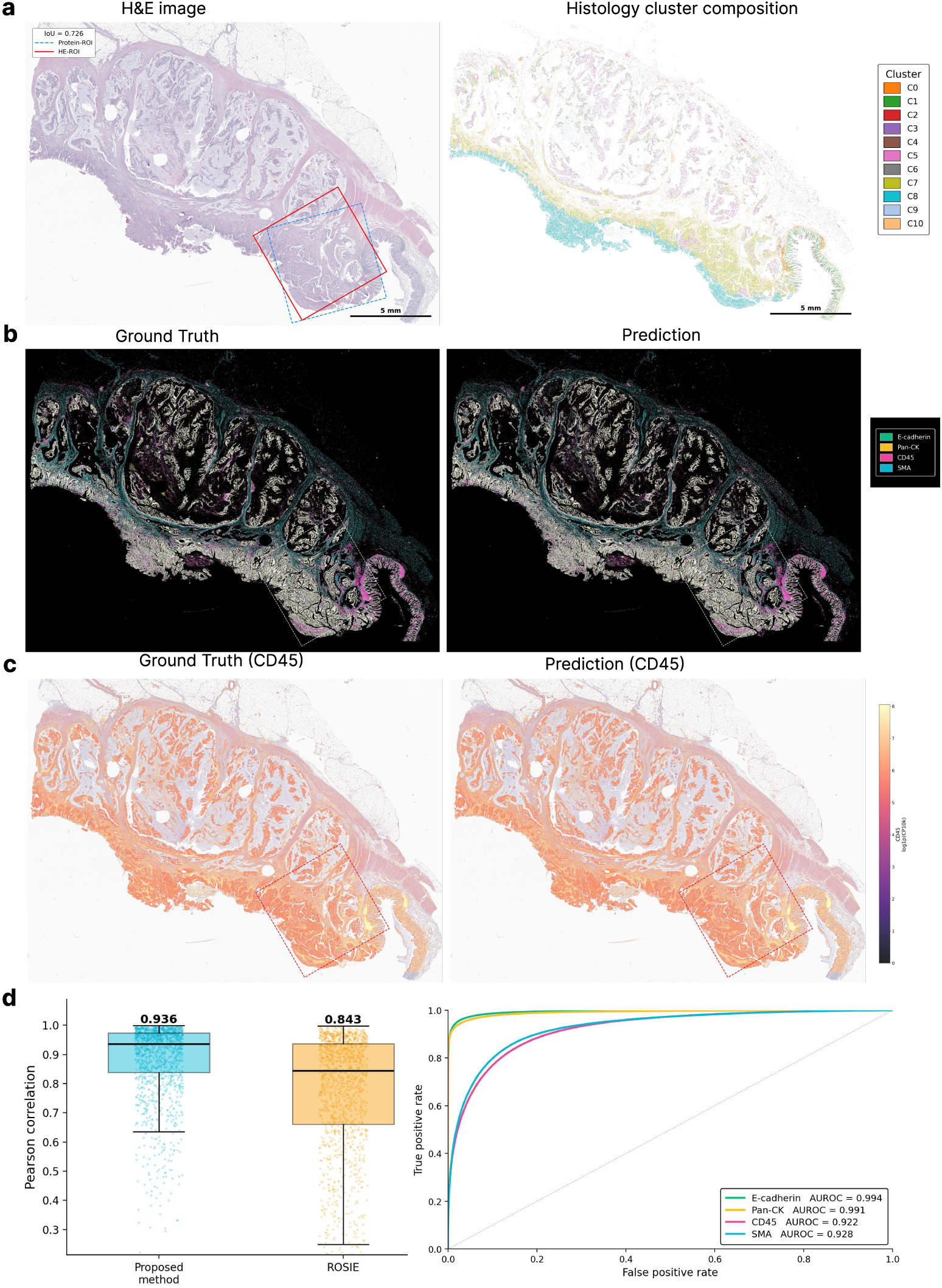
RECON infers ROIs and reconstructs spatial proteomics at single-cell resolution. **a**, Orion colorectal cancer section CRC01. Left: H&E image with the H&E-inferred ROI (HE-ROI) and the ROI obtained from measured protein expression (Protein-ROI). Right: histological cluster map used for ROI scoring. **b**, Measured and reconstructed expression of four representative markers, shown as a composite pseudo-immunofluorescence image. **c**, Measured and reconstructed CD45 intensity, overlaid on the H&E section. The dashed outline marks the ROI. **d**, Left: per-cell Pearson correlation between measured and reconstructed expression across all 16 markers for RECON and ROSIE, computed on cells outside the ROI. Right: ROC curves for marker positivity after gating, with the area under the curve given for each marker.

In addition, RECON achieved higher single-cell reconstruction accuracy than GHIST [27] on both platforms datasets (Fig. 3c). for the V1 section, the per-cell Pearson correlation between predicted and measured expression was 0.633 for RECON versus 0.578 for GHIST, while for the Prime section, the corresponding correlations were 0.240 versus 0.127, respectively.

If reconstruction primarily reflected memorization of the training data, the model performance would be expected to decline sharply with increasing distance from the ROI. Instead, the accuracy degraded gradually with spatial distance, supporting model generalization beyond the training region. In the V1 data, the per-cell Pearson correlation was 0.708 immediately adjacent to the ROI and gradually decreased to approximately 0.583 (Supplementary Fig. 3b). The Prime data showed a similar gradual trend (from 0.239 to 0.208; Supplementary Fig. 4b).

Together, these results demonstrate the utility of RECON for spatial transcriptomics: (1) H&E-based ROI selection identified regions as informative as those selected using molecular data, achieving comparable reconstruction performance; (2) single-cell expression reconstructed from the selected ROI achieved higher accuracy than the cell-level baseline across two Xenium platforms; and (3) reconstruction accuracy also extended to regions far from the ROI.

### RECON infers ROIs and reconstructs spatial proteomics at single-cell resolution

Because the framework makes no assumptions specific to transcriptomic data, we next asked whether RECON could be applied directly to other molecular modalities. We applied RECON to spatial proteomics using the Orion CRC dataset [3]. Orion acquires H&E and mIF images from the same physical slide, providing intrinsically aligned morphology and protein measurements at single-cell resolution and thus a registration-free reference for evaluating reconstruction accuracy.

We first evaluated RECON using the CRC01 section, comprising approximately 1.62 million cells and 16 protein markers including E-cadherin, Pan-CK, CD45 and SMA. Using the same architecture as for spatial transcriptomics, RECON inferred an ROI from the H&E images, trained on the protein readouts of the 390,000 cells within the ROI (23.9% of the section), and reconstructed single-cell protein expression for the remaining 1.23 million cells (Fig. 4).

Similarly, the H&E ROI selected by RECON was compared with the one obtained by clustering cells on their measured protein expression (Protein ROI), achieving an IoU of 0.726 (Fig. 4a). We evaluated reconstruction at two levels: continuous expression and marker gating, with all metrics computed on held-out cells outside the ROI. Reconstructed CD45 recapitulated the regional intensity distribution of the measured ground truth data (Fig. 4c), and the median per-cell Pearson correlation across markers was 0.936 for RECON versus 0.843 for ROSIE [28] (Fig. 4d). For marker gating, we binarized reconstructed and measured intensities by fitting a two-component Gaussian mixture per marker, as is standard for CyCIF data [34]. Marker-positive cells were reproduced in their correct spatial distribution (Fig. 4b), with AUROC values of 0.994, 0.991, 0.928 and 0.922 for E-cadherin, Pan-CK, SMA and CD45, respectively (Fig. 4d).

Performance was reproduced in a second, independent Orion section (CRC03): median per-cell Pearson was 0.912 for RECON versus 0.835 for ROSIE, and median per-marker Pearson correlation across the 16 markers was 0.549 for RECON versus 0.014 for ROSIE (Supplementary Fig. 5). This difference in per-marker accuracy follows from the unit of computation. Pixel-level predictions of ROSIE must be aggregated to single cells, a step that averages away between-cell variation within each marker, whereas RECON operates directly at single-cell resolution.

### RECON makes tumor immune microenvironment architecture accessible from routine H&E images

The preceding sections established that RECON reconstructs single-cell molecular profiles beyond the selected ROI. Numerical accuracy, however, does not guarantee that spatial organization is preserved: a reconstruction can be accurate per cell, yet fail to recover coherent tissue structures. We therefore asked whether RECON could recover microenvironmental features that are prognostically and immunotherapetically relevant, namely tertiary lymphoid structures, immune infiltration, and the tumor boundary. In a gastric cancer Xenium section (Fig. 5a, b), we compared RECON’s reconstruction with the measured ground truth of the same section.

**Fig. 5.**
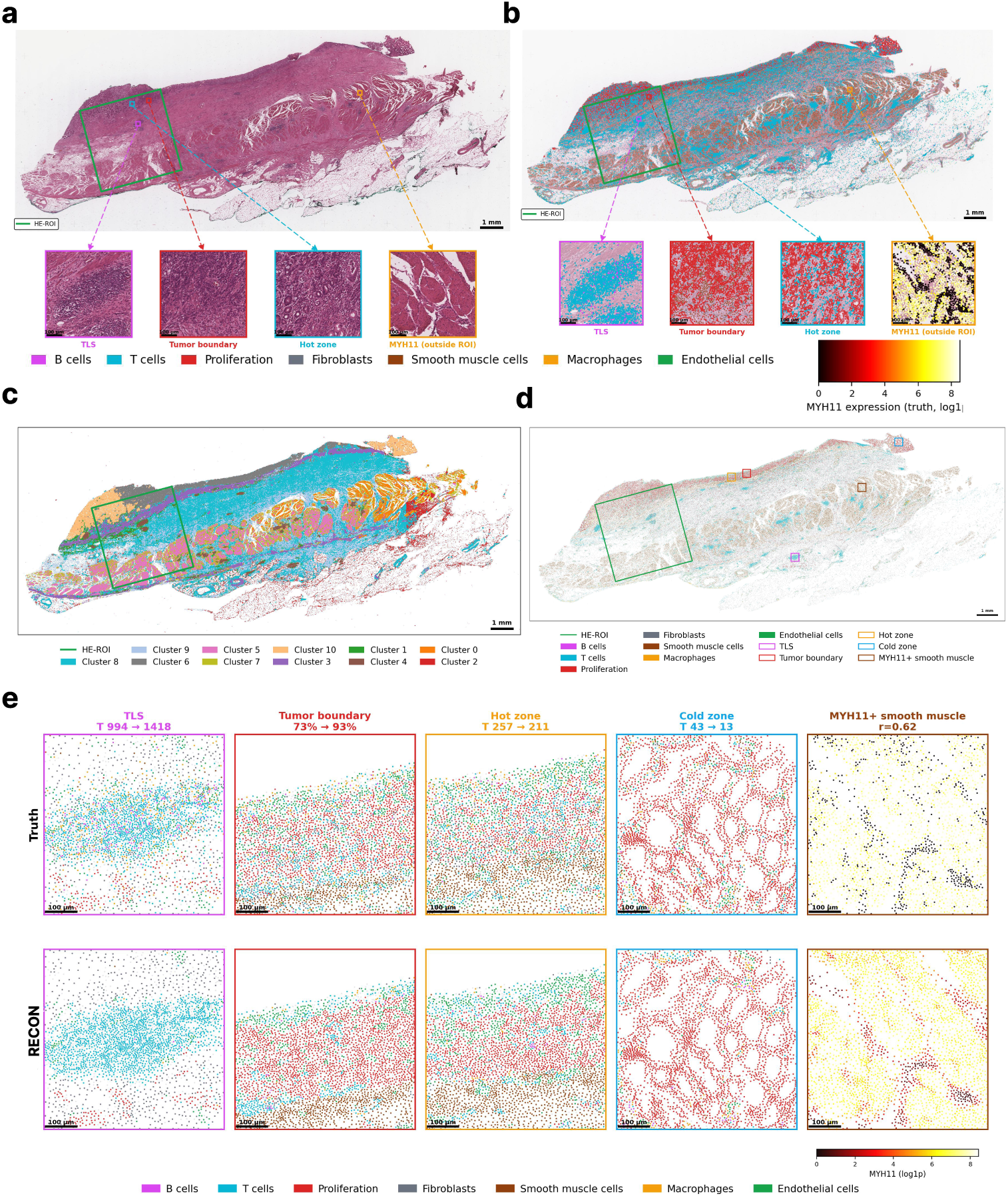
RECON recovered microenvironmental structures from routine H&E. **a**, A gastric cancer Xenium section with the inferred H&E ROI and magnified views of a tertiary lymphoid structure (TLS), the tumour boundary, an immune-dense region (hot zone), and an MYH11-positive smooth muscle region lying outside the ROI. **b**, Measured cell types and MYH11 expression at the same locations. **c**, Histological cluster map used for ROI scoring. **d**, Cell types reconstructed by RECON across the section, with the regions examined in **e** outlined. **e**, Measured (top) and reconstructed (bottom) cell types at each region. Above each panel, the number of T cells or the tumor fraction is provided for the measured and reconstructed data; for the MYH11-positive region, the Pearson correlation (*r*) between measured and reconstructed expression is given.

RECON recovered several clinically relevant structures from the reconstruction (Fig. 5c–e). The region-by-region comparison with ground truth is presented in Fig. 5e. B-cell prediction was unreliable at the single-cell level, so we used T-cell density as a proxy for lymphoid aggregation. RECON captured a T-cell-enriched aggregate at a location consistent with the measured data. The boundary between tumor parenchyma and stroma was likewise delineated, with immune-dense regions (hot zone) contrasting with sparsely infiltrated ones (cold zone). A MYH11-positive smooth muscle region was also correctly localized.

Localization, however, did not extend to quantification. Tumor proportions were overestimated in the transition zone at the boundary, immune cells were underestimated in sparsely infiltrated regions, and the cell count of the MYH11-positive region departed from the measured value. Thus, RECON captured the spatial location of these structures but was less accurate in estimating how many cells they contain.

This case nonetheless demonstrates that a single routine H&E section processed by RECON can be used to locate clinically relevant structures at a whole-slide scale. Future work will focus on improving predictions for sparse populations, such as B cells, and evaluating clinical utility in larger cohorts with paired outcome data.

### Ablation studies

Three design choices underpin RECON: single cells rather than superpixels as the computational unit, morphological features drawn from multiple scales and network depths, and output dimensionality K (Methods). We isolated the contribution of each design choice through controlled ablations in both transcriptomics and proteomics modalities, evaluating variants using Pearson correlation on cells outside the ROI (Supplementary Fig. 6).

The computational unit mattered for transcriptomics but not for proteomics (Supplementary Fig. 6a, d). Replacing single-cell features with superpixel features reduced the genewise correlation from 0.258 to 0.211 for RNA, with all three Xenium data sections shifting in the same direction, whereas the proteomic change was within expected variability (0.574 to 0.563; one section improved, one unchanged).

The richness of the morphological representation affected reconstruction accuracy both modalities (Supplementary Fig. 6b, e). Accuracy improved as features from local CLS alone were progressively augmented with contextual scales and intermediate encoder layers to the full 4,096-dimensional representation, which performed best in both modalities (transcriptome: 0.220 to 0.258; protein: 0.470 to 0.567). Morphological information from multiple spatial scales and network depths, therefore, contributes substantially to molecular reconstruction.

The dimensionality K of the output head determines the latent space in which predictions are reconstructed (Supplementary Fig. 6c, f). For the transcriptome, accuracy increased gradually with K before plateauing, whereas removing the bottleneck entirely degraded performance, indicating that a low-dimensional bottleneck acts as a regularizer. For protein, accuracy plateaued at K = 16, consistent with a 16-marker panel. We therefore adopted K = 16 for both modalities.

## Discussion

We developed RECON, a framework that unifies ROI inference with out-of-ROI molecular reconstruction from H&E images at single-cell resolution. RECON identifies the most representative ROI and, using molecular measurements acquired within that ROI for training, reconstructs continuous expression and predicts cell types for individual remaining cells on the section. The inferred ROIs agreed more closely with expert annotation than superpixel-based selection (IoU: 0.75 versus 0.64), and the resulting reconstructions outperformed the cell-level transcriptomic baseline GHIST [27] and the pixel-level proteomic baseline ROSIE [28].

The contribution of RECON lies in unifying three elements: single cells as the computational unit, molecular reconstruction from a measured ROI, and cross-modal generality. First, unlike S2-omics [7], RECON operates on segmented cells rather than superpixels and returns continuous molecular profiles rather than discrete labels, improving on both tasks addressed by S2-omics. ROIs selected by RECON agree more closely with expert annotations, particularly at small ROI sizes, where superpixel-based selection can fall beyond the profiled field of view. Working at single-cell resolution also yields a molecular profile and a cell type for every cell, which is required whole-slide structural analysis. Second, RECON enables extrapolation from a single measured ROI. RECON is designed around the practical constraint that only one ROI per section is typically assayed and predicts from measurements within that region rather than relying on whole-section supervision or multi-section integration. Third, RECON demonstrates cross-modal generality. A single architecture handled gene panels from 313 to 5,000 targets and a 16-marker protein panel, requiring no architectural changes beyond the output dimension, while outperforming the cell-level baseline GHIST [27] and the pixel-level baseline ROSIE [28], respectively. These results suggest that RECON learns a general mapping from cellular morphology to molecular readout rather than a modality-specific fit.

Spatial omics assays remain expensive and have limited throughput. RECON makes it feasible to profile only a small region and recover whole-section molecular information from routine H&E images. Two observations suggest that this recovery is based on genuine morphological signals rather than on proximity to the training data. An ROI selected from histology substantially overlapped one chosen using measured expression data, and the quality of reconstruction did not depend on the precise placement of the ROI. On this basis, RECON delineated tertiary lymphoid structures, tumor boundaries, and regions of differing immune density directly from H&E images. Although a pathologist can recognize such features morphologically, RECON assigns them a single-cell molecular composition, bringing quantitative interpretation of the tumor immune microenvironment into the conventional histology workflow.

Several limitations should be considered when interpreting these conclusions. First, the accuracy gain from single-cell granularity is modality-dependent: substantial for transcriptomics, modest for proteomics. Second, evaluation of ROI inference relies on expert annotation, which is itself subjective. Agreement with a pathologist therefore demonstrates plausibility rather than optimality, and whether a computationally selected ROI yields more biological insight than a manually drawn one remains untested. Third, reconstruction recovers the spatial location of structures more reliably than it quantifies their cellular composition, and rare populations such as B cells are poorly predicted, limiting direct identification of tertiary lymphoid structures. Fourth, the analysis of microenvironmental structures is a single-section case study, without systematic validation in larger cohorts with paired clinical outcomes.

These limitations provide the direction for future work. Improving prediction for rare cell types, particularly B cells, would strengthen the reliability of identifying immune structures such as tertiary lymphoid structures. Evaluation in larger, multi-cancer cohorts, together with analyses linking recovered structures to patient outcomes such as treatment response and prognosis, is necessary before structural recovery can translate into clinical utility. Methodologically, RECON’s prediction head is a minimal multilayer perceptron, allowing the same architecture to transfer across modalities while leaving cell-cell spatial context encoded only implicitly in the input features, making its explicit modeling a natural extension. Although RECON is designed to reconstruct whole-slide molecular profiles from a single measured ROI, the framework can also be extended to incorporate multiple ROIs within a tissue section. As paired H&E and spatial omics datasets accumulate, the question shifts from whether morphology predicts molecular state to how much of the molecular landscape is morphologically determined. Answering this question would establish inexpensive histology as a scalable route connecting routine pathology to spatial biology.

## Methods

Figure 1 summarizes the framework of RECON. RECON uses two adjacent sections from the same FFPE tissue block: one imaged by H&E and the other profiled by spatial omics (Fig. 1a). In Stage 1, RECON selects an ROI from the H&E section (Fig. 1b). In Stage 2, RECON trains deep learning models based on measurements acquired within ROI and reconstructs molecular profiles beyond the measured ROI (Fig. 1c), yielding a continuous expression profile and a cell type for each cell across the slide (Fig. 1d). Both stages share a frozen UNI backbone and operate without modification on spatial transcriptomics and spatial proteomics. Each component is described in detail below.

### Cell segmentation and feature extraction

The two stages of RECON use different cell segmentation strategies tailored to their respective task settings. Stage 1 (ROI selection) relies entirely on H&E images and uses no molecular measurements; therefore, cells are segmented from H&E with CellViT++ across all datasets (Fig. 1b). This reproduces the intended application scenario, in which segmentation and ROI selection are carried out from the H&E alone, before any molecular measurement is performed. Stage 2 (molecular reconstruction) uses the native segmentation from each spatial omics platform (Xenium and Orion respectively), so that the predictions and the ground truth are evaluated under a single cell definition. The two stages are linked through the selected rectangular ROI. Because the ROI is a geometric region rather than a set of cells, it transfers between the two segments without requiring cell-level correspondence: natively segmented cells falling inside the ROI provide the supervisory signal for Stage 2, and those outside serve as prediction targets.

Let *c_i_* denote the centroid of a cell *i*. From the H&E image, a local patch (64 *×* 64) and a context patch (224 *×* 224) were cropped centered on *c_i_*, both resized to 224 *×* 224 and denoted as *x_i_*^loc^ and *x_i_*^ctx^. Embeddings were then extracted using a frozen UNI backbone *f*_UNI_.

### Stage 1: ROI selection

Stage 1 of RECON infers the ROI in four sequential steps: cell feature learning, grid assignment, clustering, and ROI scoring.

#### Cell feature learning

UNI [30] returns a class (CLS) token that aggregates information across the whole patch through self-attention and serves as a holistic patch representation. For each cell, we encoded both image patches (*x_i_*^loc^ and *x_i_*^ctx^) with UNI and *ℓ*_2_-normalized the resulting CLS tokens separately, as given in Equation (1):

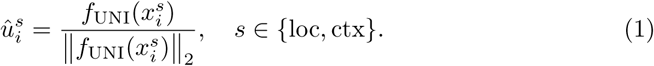

The two normalized 1024-dimensional tokens were concatenated and subsequently normalized to obtain the Stage 1 cell feature *z_i_*, as given in Equation (2):

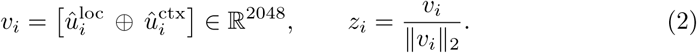

where ⊕ denotes vector concatenation. Together, cells are represented as 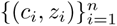, where *n* is the number of cells, *c_i_* is the centroid of a cell *i*, and *z_i_* is the cell feature.

#### Grid assignment

Because cells are distributed at irregular spatial coordinates, we mapped them onto an 8 *µ*m grid for downstream tasks. For each bin *b*, we retained the feature of the cell closest to the bin center rather than averaging features over the cells within the bin, as given in Equation (3):

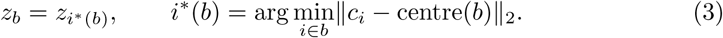

Bins containing no cell were left empty and excluded from clustering. Thus, each occupied bin retains the feature of a single cell, preserving single-cell granularity while providing a regular spatial representation. Our approach differs from superpixel methods, which aggregate over multiple cells within each spatial unit.

#### Clustering

Bin-level features were reduced by Principal Component Analysis (PCA) to 128 components, retaining approximately 65% of the variance. The groups were then partitioned into 20 groups by *k*-means with *k*-means++ initialization, and merged to *k^′^* histological clusters by average-linkage hierarchical clustering on the cluster centroids, following the merge over clusters procedure of S2-omics [7].

#### ROI scoring and selection

ROI scoring and selection follow the rectangular ROI algorithm of S2-omics [7], with superpixels replaced by the single-cell cluster map described above. Each candidate rectangle is scored by the weighted geometric mean of three components: *coverage*, the fraction of quality-control passing units within the ROI; *balance*, the cosine similarity between the ROI’s cluster distribution and a uniform distribution over the *k^′^* histological clusters; and *size*, a logistic transform of the effective sampling rate, as given in Equation (4):

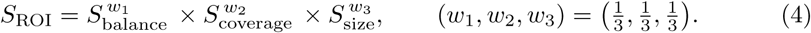

Candidate rectangles were sampled across the slide following the default settings of S2-omics. The highest-scoring rectangle was retained as the selected ROI, and the cells *A* within it provided the supervisory signal for Stage 2.

### Stage 2: Molecular reconstruction

#### Input features

Two complementary readouts were taken from the frozen encoder for both image patches (*x_i_*^loc^ and *x_i_*^ctx^): the class token cls(*·*), which summarizes the patch as a whole, and the central patch token ctr(*·*), which corresponds to the 16 *×* 16 region at the patch center and therefore provides a more localized representation of the cell (*x_i_*^loc^ and *x_i_*^ctx^), RECON extracts both at the local and the context scale, as given in Equation (5) (Fig. 1c):

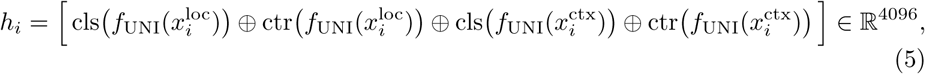

where *⊕* denotes concatenation, and each component was layer-normalized beforehand. The 4096-dimensional *h_i_* was reduced to 128 dimensions by PCA, giving *g_i_ ∈* R^128^, which served as the shared input to two separately trained prediction branches.

#### Regression predictor (gene or protein expression)

Measured expression was normalized as log_1*p*_(CP10k) and denoted *y_i_ ∈* R*^G^*, where *G* is the number of genes or proteins. An output-side PCA, *P* (*·*), was fitted on the cells within the ROI (*A*, hereafter anchor cells) alone, compressing expression into a *K*-dimensional latent space that served as the regression target *t_i_* = *P* (*y_i_*) (*K* = 16 for transcriptomics; for the 16-marker protein panel, *K* = 16 is equivalent to direct regression). The regressor *ϕ*_reg_ is a three-layer MLP [Linear(128 *→* 256) *→* ReLU *→* Dropout(0.1) *→* Linear(256 *→* 256) *→* ReLU *→* Dropout(0.1) *→* Linear(256 *→ K*)], trained on the anchor cells with a smooth *L*_1_ (Huber) loss, as given in Equation (6):

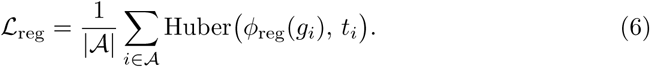

At inference, *t*^^^*_i_* was predicted for each cell outside the ROI and mapped back to the original expression space by the inverse transform, *y*^*_i_* = *P^−^*^1^(*t*^^^*_i_*).

### Classification predictor (cell type)

Cell-type labels *ℓ_i_ ∈ {*1*, &, N}* were defined by marker-based annotation of the anchor cells, giving *N* classes. The classifier *ϕ*_cls_ is a separate three-layer MLP (identical in architecture to *ϕ*_reg_, with an *N* -dimensional output layer), trained on the anchor cells with a cross-entropy loss, as given in Equation (7):

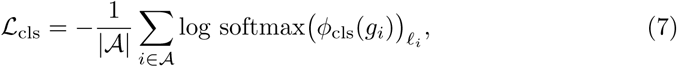

where softmax(*·*)*_ℓi_* is the probability assigned by the predicted distribution to the true class *ℓ_i_*. At inference, cell types were predicted for every cell outside the ROI.

#### Training

The two predictors were trained independently using soley the anchor cells. Both predictors were optimized with AdamW (learning rate: 1 *×* 10*^−^*^3^; weight decay: 1 *×* 10*^−^*^5^), a batch size of 512, and 20 epochs.

### Evaluation

All metrics were computed on cells outside the selected ROI. Let *Y, Y*^^^ *∈* R*^M×G^* be the measured and predicted expression over these *M* cells and *G* features (genes or protein markers). We reported two Pearson correlations, as given in Equation (8): *r*_cell_ across features within each cell and *r*_gene_ across cells for each feature.

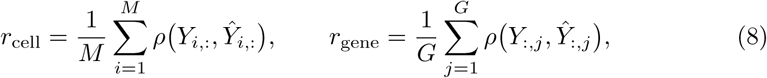

where *ρ* is the Pearson correlation coefficient.

## Data availability

We analyzed the following datasets: (1) 10x Genomics human colorectal cancer data, sample P1CRC, with Visium HD and Xenium In Situ performed on serial sections from the same FFPE block (ref. 31; https://www.10xgenomics.com/platforms/visium/product-family/dataset-human-crc); (2) 10x Genomics human breast cancer Xenium data, in situ sample 1, replicate 1 (denoted Xenium S1; ref. 33; https://www.10xgenomics.com/products/xenium-in-situ/preview-dataset-human-breast); (3) 10x Genomics Xenium Prime FFPE human breast cancer data, acquired with the 5K Human Pan Tissue and Pathways panel plus 100 custom genes (denoted Xenium Prime; https://www.10xgenomics.com/datasets/xenium-prime-ffpe-human-breast-cancer); (4) human colorectal cancer Orion high-plex immunofluorescence and H&E data generated by Lin et al. (ref. 3; samples CRC01 and CRC03), available via Zenodo at https://zenodo.org/records/7637988); (5) human gastric cancer Xenium data generated by the Tae Hyun Hwang laboratory and available via Zenodo at https://zenodo.org/records/15164980) (denoted Xenium 15164980).

## Code availability

The source code for RECON is available at https://github.com/cwanlab/RECON.

## Acknowledgements

G.W. is supported in part by the National Cancer Institute of the National Institutes of Health under Award Number R00CA286966.

## Author contributions

X.Y. and G.W. conceptualized and designed the project. X.Y. implemented the RECON framework and conducted the experiments. X.Y., N.H., and G.W. drafted the manuscript. C.G.L. and A.R. provided pathological expertise. G.W. provided project resources and supervision. All authors reviewed and approved the final manuscript.

## References

[1] Janesick, A., Shelansky, R., Gottscho, A.D., Wagner, F., Williams, S.R., Rouault, M., Beliakoff, G., Morrison, C.A., Oliveira, M.F., Sicherman, J.T., et al.: High resolution mapping of the tumor microenvironment using integrated single-cell, spatial and in situ analysis. Nature communications 14(1), 8353 (2023)

[2] Goltsev, Y., Samusik, N., Kennedy-Darling, J., Bhate, S., Hale, M., Vazquez, G., Black, S., Nolan, G.P.: Deep profiling of mouse splenic architecture with codex multiplexed imaging. Cell 174(4), 968–981 (2018)

[3] Lin, J.-R., Chen, Y.-A., Campton, D., Cooper, J., Coy, S., Yapp, C., Tefft, J.B., McCarty, E., Ligon, K.L., Rodig, S.J., et al.: High-plex immunofluorescence imaging and traditional histology of the same tissue section for discovering image-based biomarkers. Nature cancer 4(7), 1036–1052 (2023)

[4] Elhanani, O., Ben-Uri, R., Keren, L.: Spatial profiling technologies illuminate the tumor microenvironment. Cancer cell 41(3), 404–420 (2023)

[5] De Visser, K.E., Joyce, J.A.: The evolving tumor microenvironment: From cancer initiation to metastatic outgrowth. Cancer cell 41(3), 374–403 (2023)

[6] Hallinan, C., Ji, H.J., Tsou, E., Salzberg, S.L., Fan, J.: Evidence of off-target probe binding affecting 10x genomics xenium gene panels compromise accuracy of spatial transcriptomic profiling. Elife 14, 107070 (2026)

[7] Yuan, M., Jin, K., Yan, H., Schroeder, A., Luo, C., Yao, S., Dumoulin, B., Levinsohn, J., Luo, T., Clemenceau, J.R., et al.: Smart spatial omics (s2-omics) optimizes region of interest selection to capture molecular heterogeneity in diverse tissues. Nature cell biology, 1–14 (2025)

[8] Gindra, R.H., Palla, G., Nguyen, M., Wagner, S.J., Tran, M., Theis, F.J., Saur, D., Crawford, L., Peng, T.: A large-scale benchmark of cross-modal learning for histology and gene expression in spatial transcriptomics. In: Proceedings of the IEEE/CVF International Conference on Computer Vision, pp. 1182–1192 (2025)

[9] Wang, C., Chan, A.S., Fu, X., Ghazanfar, S., Kim, J., Patrick, E., Yang, J.Y.: Benchmarking the translational potential of spatial gene expression prediction from histology. Nature Communications 16(1), 1544 (2025)

[10] Zhang, D., Schroeder, A., Yan, H., Yang, H., Hu, J., Lee, M.Y., Cho, K.S., Susztak, K., Xu, G.X., Feldman, M.D., et al.: Inferring super-resolution tissue architecture by integrating spatial transcriptomics with histology. Nature biotechnology 42(9), 1372–1377 (2024)

[11] He, B., Bergenstråhle, L., Stenbeck, L., Abid, A., Andersson, A., Borg, Å., Maaskola, J., Lundeberg, J., Zou, J.: Integrating spatial gene expression and breast tumour morphology via deep learning. Nature biomedical engineering 4(8), 827–834 (2020)

[12] Polański, K., Bartolomé-Casado, R., Sarropoulos, I., Xu, C., England, N., Jahnsen, F.L., Teichmann, S.A., Yayon, N.: Bin2cell reconstructs cells from high resolution visium hd data. Bioinformatics 40(9), 546 (2024)

[13] Kamel, M., Song, Y., Solbas, A., Villordo, S., Sarangi, A., Senin, P., Sunaal, M., Ayestas, L.C., Levin, C., Wang, S., et al.: Enact: end-to-end analysis of visium high definition (hd) data. Bioinformatics 41(3), 094 (2025)

[14] Stringer, C., Wang, T., Michaelos, M., Pachitariu, M.: Cellpose: a generalist algorithm for cellular segmentation. Nature methods 18(1), 100–106 (2021)

[15] Hörst, F., Rempe, M., Heine, L., Seibold, C., Keyl, J., Baldini, G., Ugurel, S., Siveke, J., Grünwald, B., Egger, J., et al.: Cellvit: Vision transformers for precise cell segmentation and classification. Medical image analysis 94, 103143 (2024)

[16] Hörst, F., Rempe, M., Becker, H., Heine, L., Keyl, J., Kleesiek, J.: Cellvit++: Energy-efficient and adaptive cell segmentation and classification using foundation models. Computer Methods and Programs in Biomedicine, 109206 (2026)

[17] Graham, S., Vu, Q.D., Raza, S.E.A., Azam, A., Tsang, Y.W., Kwak, J.T., Rajpoot, N.: Hover-net: Simultaneous segmentation and classification of nuclei in multi-tissue histology images. Medical image analysis 58, 101563 (2019)

[18] Zhang, P., Gao, C., Zhang, Z., Yuan, Z., Zhang, Q., Zhang, P., Du, S., Zhou, W., Li, Y., Li, S.: Systematic inference of super-resolution cell spatial profiles from histology images. Nature Communications 16(1), 1838 (2025)

[19] Wu, Y., Zhou, J.-Y., Yao, B., Cui, G., Zhao, Y.-L., Gao, C.-C., Yang, Y., Zhang, S., Yang, Y.-G.: Stascan deciphers fine-resolution cell distribution maps in spatial transcriptomics by deep learning. Genome Biology 25(1), 278 (2024)

[20] Zhao, W., Liang, Z., Huang, X., Huang, Y., Yu, L.: Hist2cell: deciphering fine-grained cellular architectures from histology images. Cell Genomics 6(3) (2026)

[21] Chung, Y., Ha, J.H., Im, K.C., Lee, J.S.: Accurate spatial gene expression prediction by integrating multi-resolution features. In: Proceedings of the IEEE/CVF Conference on Computer Vision and Pattern Recognition, pp. 11591–11600 (2024)

[22] Schmauch, B., Romagnoni, A., Pronier, E., Saillard, C., Maillé, P., Calderaro, J., Kamoun, A., Sefta, M., Toldo, S., Zaslavskiy, M., et al.: A deep learning model to predict rna-seq expression of tumours from whole slide images. Nature communications 11(1), 3877 (2020)

[23] Li, Z., Li, Y., Xiang, J., Wang, X., Yang, S., Zhang, X., Eweje, F., Chen, Y., Luo, X., Li, Y., et al.: Ai-enabled virtual spatial proteomics from histopathology for interpretable biomarker discovery in lung cancer. Nature Medicine, 1–14 (2026)

[24] Andani, S., Chen, B., Ficek-Pascual, J., Heinke, S., Casanova, R., Hild, B.F., Sobottka, B., Bodenmiller, B., Koelzer, V.H., et al.: Histopathology-based protein multiplex generation using deep learning. Nature Machine Intelligence 7(8), 1292– 1307 (2025)

[25] Hao, N., Yang, X., Yan, B., Li, D., Huang, J., Wu, X., Ruiz, E.S., Luzuriaga, A., Zhao, C., Wan, G.: Histopathology-centered computational evolution of spatial omics: integration, mapping, and foundation models. Briefings in Bioinformatics 27(4), 387 (2026)

[26] Schroeder, A., Loth, M.L., Luo, C., Yao, S., Yan, H., Zhang, D., Piya, S., Plowey, E., Hu, W., Clemenceau, J.R., et al.: Scaling up spatial transcriptomics for large-sized tissues: uncovering cellular-level tissue architecture beyond conventional platforms with iscale. Nature methods 22(9), 1911–1922 (2025)

[27] Fu, X., Cao, Y., Bian, B., Wang, C., Graham, D., Pathmanathan, N., Patrick, E., Kim, J., Yang, J.Y.H.: Spatial gene expression at single-cell resolution from histology using deep learning with ghist. Nature methods 22(9), 1900–1910 (2025)

[28] Wu, E., Bieniosek, M., Wu, Z., Thakkar, N., Charville, G.W., Makky, A., Schürch, C.M., Huyghe, J.R., Peters, U., Li, C.I., et al.: Rosie: Ai generation of multiplex immunofluorescence staining from histopathology images. Nature Communications 16(1), 7633 (2025)

[29] Valanarasu, J.M.J., Xu, H., Usuyama, N., Kim, C., Wong, C., Argaw, P., Shimol, R.B., Crabtree, A., Matlock, K., Bartlett, A.Q., et al.: Multimodal ai generates virtual population for tumor microenvironment modeling. Cell 189(2), 386–400 (2026)

[30] Chen, R.J., Ding, T., Lu, M.Y., Williamson, D.F., Jaume, G., Song, A.H., Chen, B., Zhang, A., Shao, D., Shaban, M., et al.: Towards a general-purpose foundation model for computational pathology. Nature medicine 30(3), 850–862 (2024)

[31] Oliveira, M.F.d., Romero, J.P., Chung, M., Williams, S.R., Gottscho, A.D., Gupta, A., Pilipauskas, S.E., Mohabbat, S., Raman, N., Sukovich, D.J., et al.: High-definition spatial transcriptomic profiling of immune cell populations in colorectal cancer. Nature genetics 57(6), 1512–1523 (2025)

[32] Lowe, D.G.: Distinctive image features from scale-invariant keypoints. International journal of computer vision 60(2), 91–110 (2004)

[33] Janesick, A., Shelansky, R., Gottscho, A.D., Wagner, F., Williams, S.R., Rouault, M., Beliakoff, G., Morrison, C.A., Oliveira, M.F., Sicherman, J.T., et al.: High resolution mapping of the tumor microenvironment using integrated single-cell, spatial and in situ analysis. Nature communications 14(1), 8353 (2023)

[34] Lin, J.-R., Izar, B., Wang, S., Yapp, C., Mei, S., Shah, P.M., Santagata, S., Sorger, P.K.: Highly multiplexed immunofluorescence imaging of human tissues and tumors using t-cycif and conventional optical microscopes. elife 7, 31657 (2018)

