## supplemental file for "RECON infers regions of interest from H&E images and reconstructs whole-slide molecular profiles at single-cell resolution"

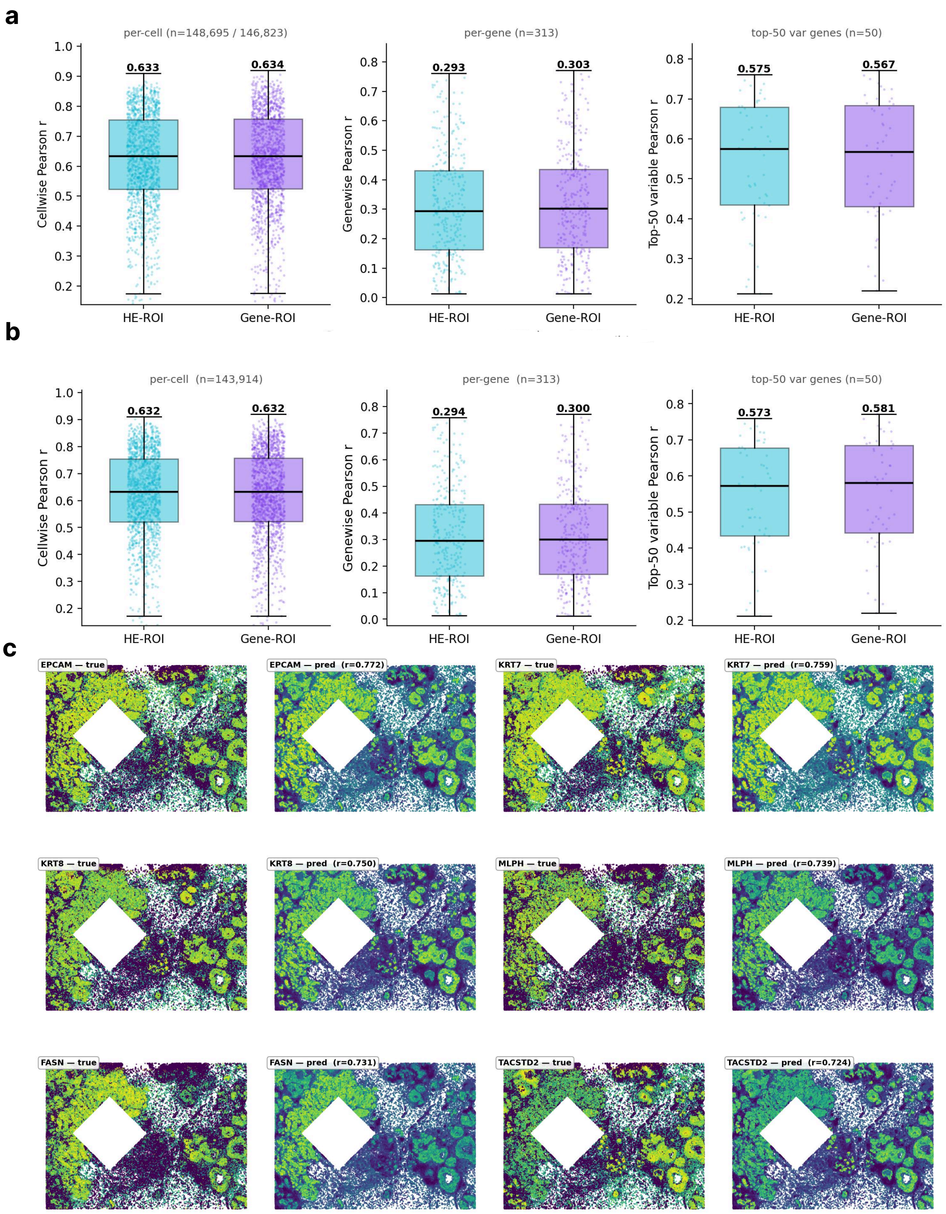


**Supplementary Figure 1 Reconstruction is insensitive to whether the ROI is chosen from histology or from measured expression (Xenium V1).** **a**, Pearson correlations (per-cell, per-gene, and top-50 variable genes) for models trained on cells within the H&E ROI and the Gene ROI, respectively, each evaluated on the cells outside its corresponding ROI (148,695 cells outside the H&E ROI and 146,823 cells outside the Gene ROI). **b**, Similar to **a**, but evaluated on the 143,914 cells lying outside both ROIs, so that the two models were compared on an identical set of cells. **c**, Measured and reconstructed expressions for the six best-predicted genes, with the selected ROI masked in white. Pearson correlation is given for each gene.

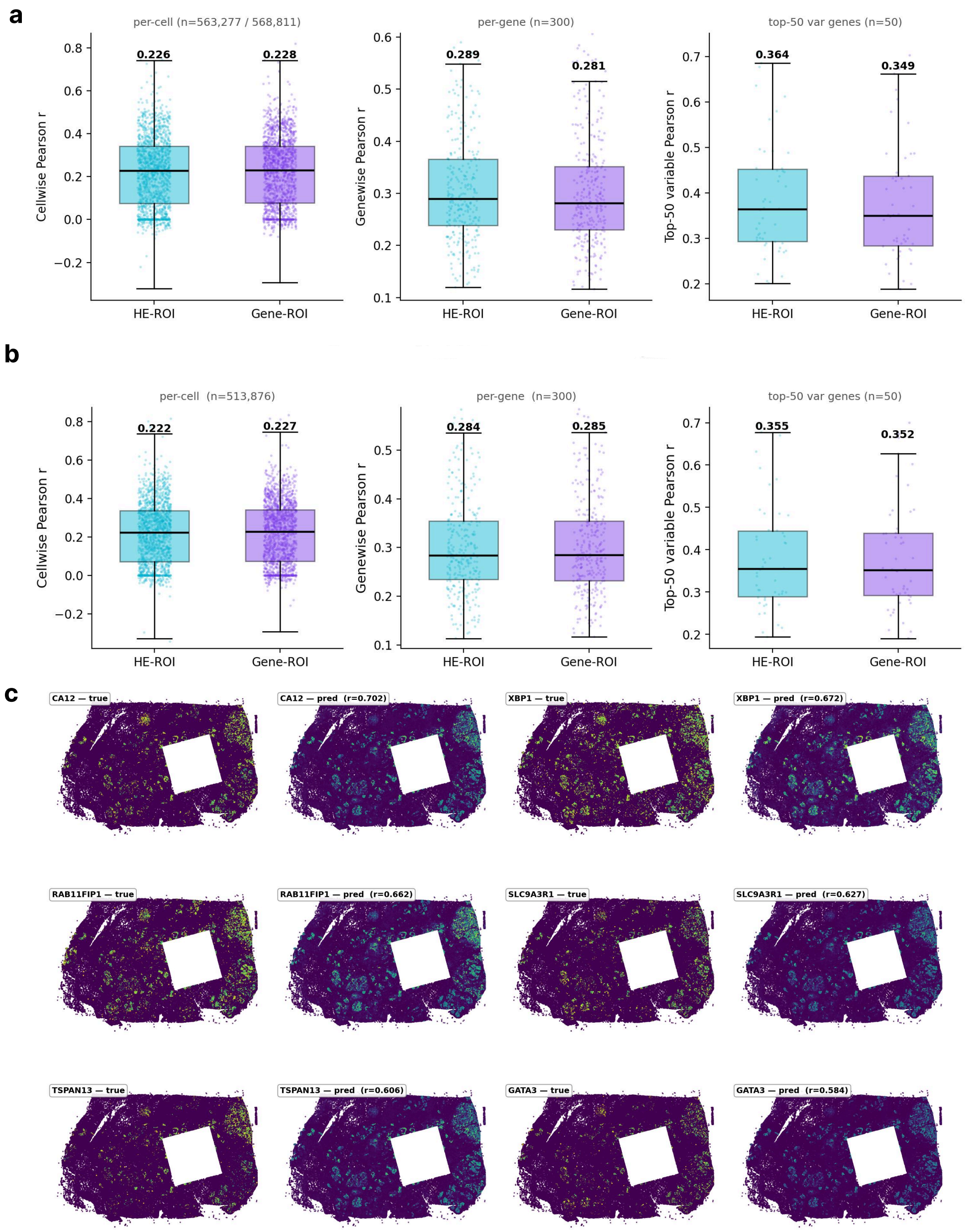


**Supplementary Figure 2 Reconstruction is insensitive to whether the ROI is chosen from histology or from measured expression (Xenium Prime).** **a**, Pearson correlations (per-cell, per-gene, and top-50 variable genes) for models trained on cells within the H&E ROI and the Gene ROI, respectively, each evaluated on the cells outside its corresponding ROI (563,277 cells outside the H&E ROI and 568,811 cells outside the Gene ROI). **b**, Similar to **a**, but evaluated on the 513,876 cells lying outside both ROIs, so that the two models were compared on an identical set of cells. **c**, Measured and reconstructed expressions for the six best-predicted genes, with the selected ROI masked in white. Pearson correlation is given for each gene.

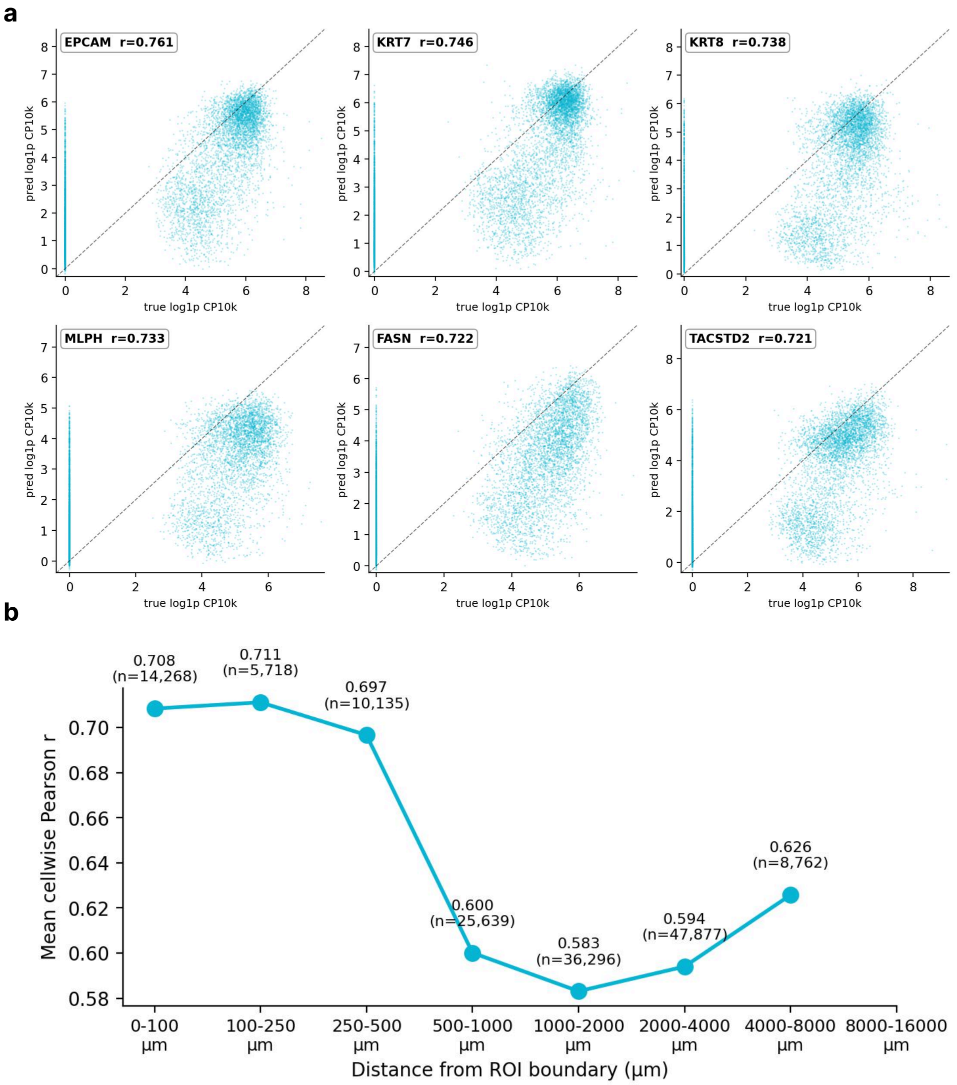


**Supplementary Figure 3 Reconstruction accuracy at the gene level and as a function of distance from the ROI. a**, Measured versus reconstructed expression for the six best-predicted genes on the Xenium V1 breast cancer section. Each point corresponds to a held-out cell. Pearson correlation is given for each gene; dashed lines indicate identity. **b**, Mean per-cell Pearson correlation as a function of distance from the ROI boundary, computed on held-out cells binned by distance. The number of cells in each bin is provided above the corresponding point.

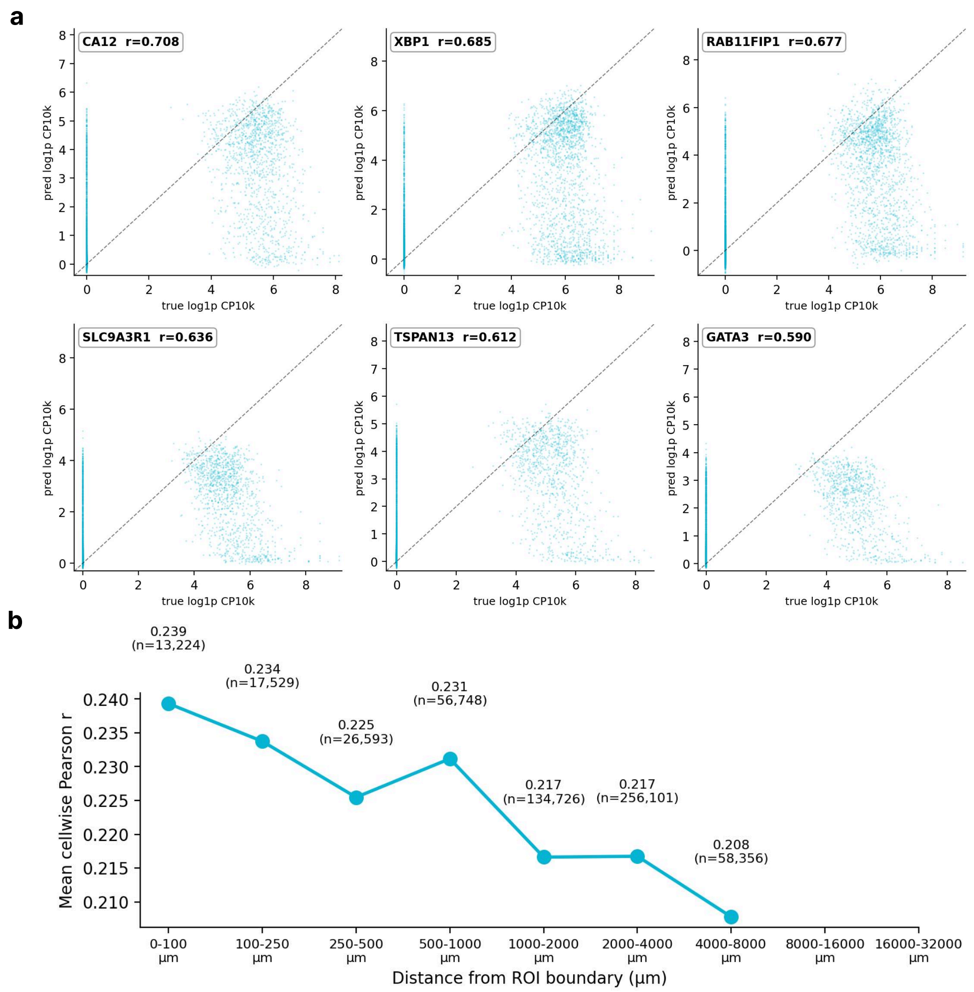


**Supplementary Figure 4 Reconstruction accuracy at the gene level and as a function of distance from the ROI (Xenium Prime).** **a**, Measured versus reconstructed expression for the six best-predicted genes. Each point corresponds to a held-out cell. Pearson correlation is provided for each gene; dashed lines indicate identity. **b**, Mean per-cell Pearson correlation as a function of distance from the ROI boundary.

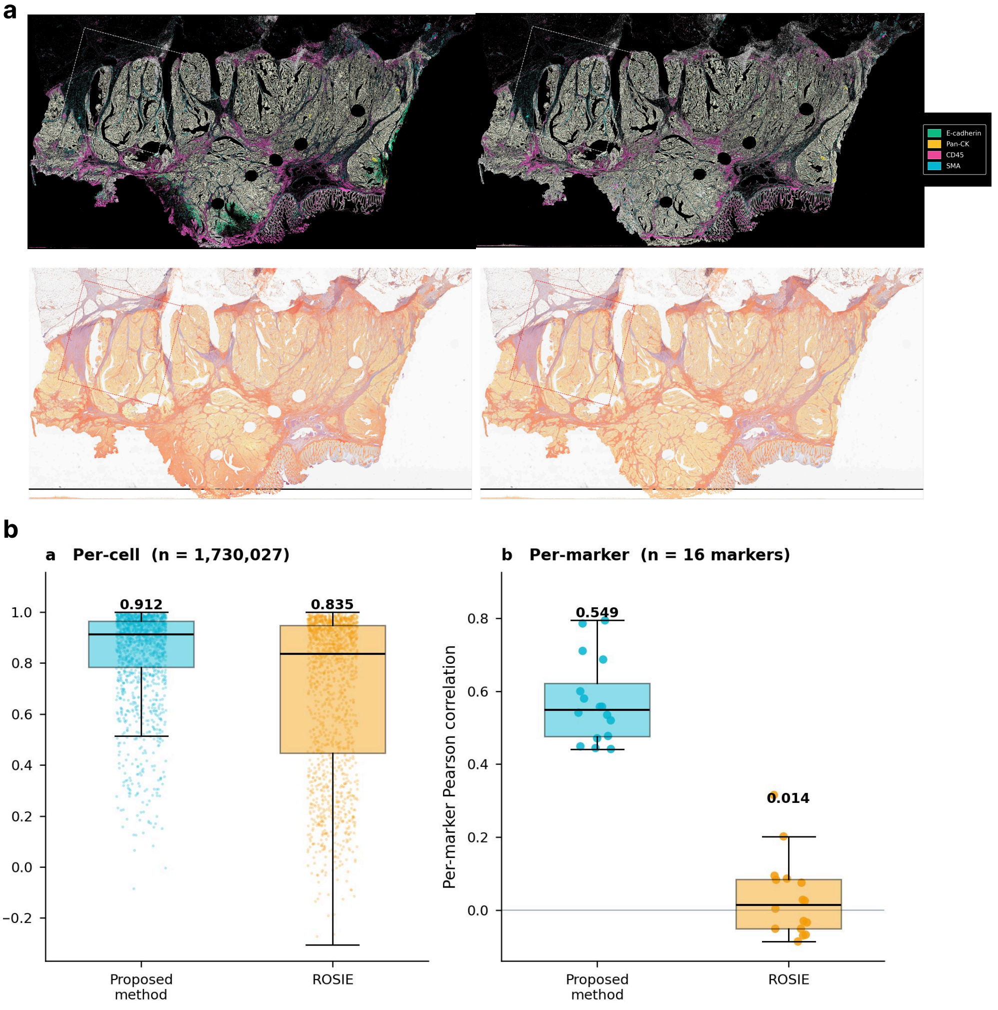


**Supplementary Figure 5 Reconstruction of single-cell protein expression on an independent Orion section (CRC03). a**, Measured (left) and reconstructed (right) protein expression. Top: composite of four representative markers. Bottom: the same section overlaid on H&E; the outline marks the selected ROI. **b**, Mean per-cell Pearson correlation across all 16 markers (left) and per-marker Pearson correlation across cells (right) for RECON and ROSIE, computed on the 1,730,027 held-out cells outside the ROI.

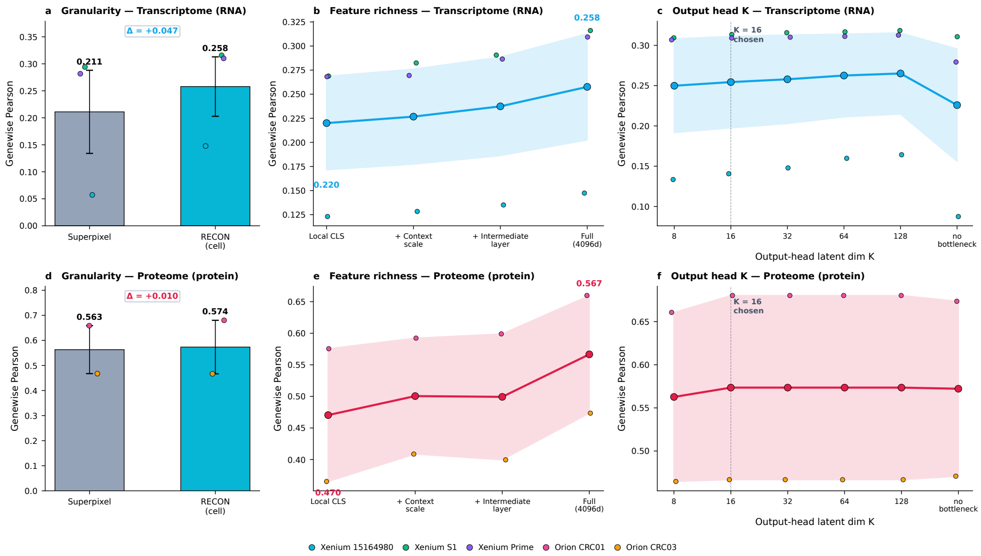


**Supplementary Figure 6 Ablation of the three design choices in RECON.** Each panel reports genewise Pearson correlation on cells outside the ROI, with one point per dataset (colours, bottom legend) and the line or bar showing the mean across datasets. **a**, **d**, Computational unit: superpixel features at matched resolution versus segmented cells, for transcriptomics and proteomics respectively. Error bars show the range across datasets. **b**, **e**, Feature richness: local CLS token alone, then adding the context scale, the intermediate-layer token, and the full 4,096-dimensional representation. Shaded bands show the range across datasets. **c**, **f**, Output-head latent dimensionality $K$, from 8 to 128 and with the bottleneck removed. The dashed line marks the value used throughout.
